# Parkinsonism disrupts the population-level organization of cortical dynamics

**DOI:** 10.64898/2026.04.21.719912

**Authors:** Yuxiao Ning, Luke A. Johnson, Jing Wang, Hiba Sheheitli, Biswaranjan Mohanty, Jerrold L. Vitek

## Abstract

Parkinson’s disease (PD) is marked by impairments in voluntary movement, including prolonged movement preparation and execution, yet how parkinsonism alters neural processing to produce these deficits remains unresolved. Prior work examining M1 spiking activity in parkinsonian states has largely characterized firing-rate changes and motor representations at the level of individual neurons, with inconsistent results and limited insight into population-level organization. Here we investigated how parkinsonism reshapes the population-level organization of neural activity in M1 during movement. We simultaneously recorded large populations of neurons from M1 in two nonhuman primates performing reaching tasks before and after induction of parkinsonism with the neurotoxin 1-methyl-4-phenyl-1,2,3,6-tetrahydropyridine (MPTP). In the parkinsonian state, both preparation and reach durations were significantly prolonged. Population-level analyses revealed that parkinsonism increased the dimensionality of M1 activity during both preparation and movement and reduced the orthogonality between preparatory and movement-related subspaces. Moreover, trial-by-trial variability in reach duration was explained by the alignment of the preparatory and reach subspaces, indicating the functional role of subspace orthogonality. Together, these findings suggest that parkinsonism disrupts the population-level organization of cortical dynamics across computations, providing a population-level framework linking altered cortical dynamics to the movement-related dysfunction observed in PD.

## Introduction

A decline in motor performance is a hallmark of Parkinson’s disease (PD), typically reflected by longer movement preparation and execution times^1,2^. Clinically, these deficits present as akinesia (the difficulty to initiate voluntary movement) and bradykinesia (the pathological slowness of movement). Dysfunction of the primary motor cortex (M1) is considered to play a crucial role in mediating the impairments of voluntary movement associated with PD^3–6^, but how altered cortical activity contributes to these deficits remains unclear.

Relatively few studies have investigated single-unit activity in M1 during active movement in the parkinsonian state and the conclusions drawn from these studies have varied. Some studies have focused on the changes in neuronal firing rates in M1, reporting reduced firing rates^3,7,8^ and decreased inhibitory responses^9^, contributing to the longstanding ‘Hypoactivation’ and ‘Hyperactivation’ debate. The hypoactivation view is grounded in the classic firing rate model of basal ganglia-thalamo-cortical network^10,11^ in which loss of dopamine in the striatum in parkinsonism results in loss of thalamocortical excitatory drive due to excessive inhibition from the basal ganglia output. The hyperactivation view, however, attributes elevated M1 firing to compensatory cortical recruitment in response to impaired basal ganglia output^5,12,13^. In addition, other studies have emphasized changes in motor representations, including a loss of directional selectivity^14^ and abnormal temporal patterning of motor commands in M1^3,15,16^. Due to relatively sparse and inconsistent data, there is little consensus on how parkinsonism alters neuronal processing in M1 during movement.

One promising avenue towards addressing these limitations is through investigating the population-level organization of neuronal activity in the parkinsonian motor cortex, which reflects how activity is coordinated across neurons over time. Over the past decade, studies of the motor cortex in healthy subjects have increasingly adopted population-level approaches, showing that structured, collective activity patterns emerge during movement preparation and execution and are not apparent from heterogeneous single-unit responses^17–20^. Additionally, population-level analyses offer single-trial statistical power by leveraging shared structure across neurons^21^, to explain the pronounced behavioral variability observed in PD.

In healthy subjects, two fundamental organizational properties on population-level neuronal activity in M1 have been proposed to underlie movement preparation and generation. First, M1 activity evolves within a low-dimensional subspace during active movement, which reflects coordinated patterns of neural activity that constrain the effective degrees of freedom used for motor control (Figure 1A, B). This low dimensionality is thought to simplify downstream readout by consolidating high-dimensional neuronal activity into a compact set of latent variables that encode movement-relevant information, thereby enabling robust and efficient control of movement^17,18,22–24^. Second, preparatory and movement-related activity in M1 occupy largely orthogonal subspaces within this low-dimensional state space, providing a mechanism to segregate motor planning from execution^19,20,25^ (Figure 1C). By confining preparatory activity to dimensions that do not directly project onto movement-generating output, this orthogonal organization allows the motor system to establish an appropriate preparatory state without prematurely driving movement. Upon movement initiation, population activity transitions into an execution-related subspace that effectively drives motor output. Whether and how these principal population-level properties of M1, low dimensionality and subspace orthogonality, are altered in parkinsonism remains unclear, despite their central role in enabling efficient movement preparation and execution.

**Figure 1:**
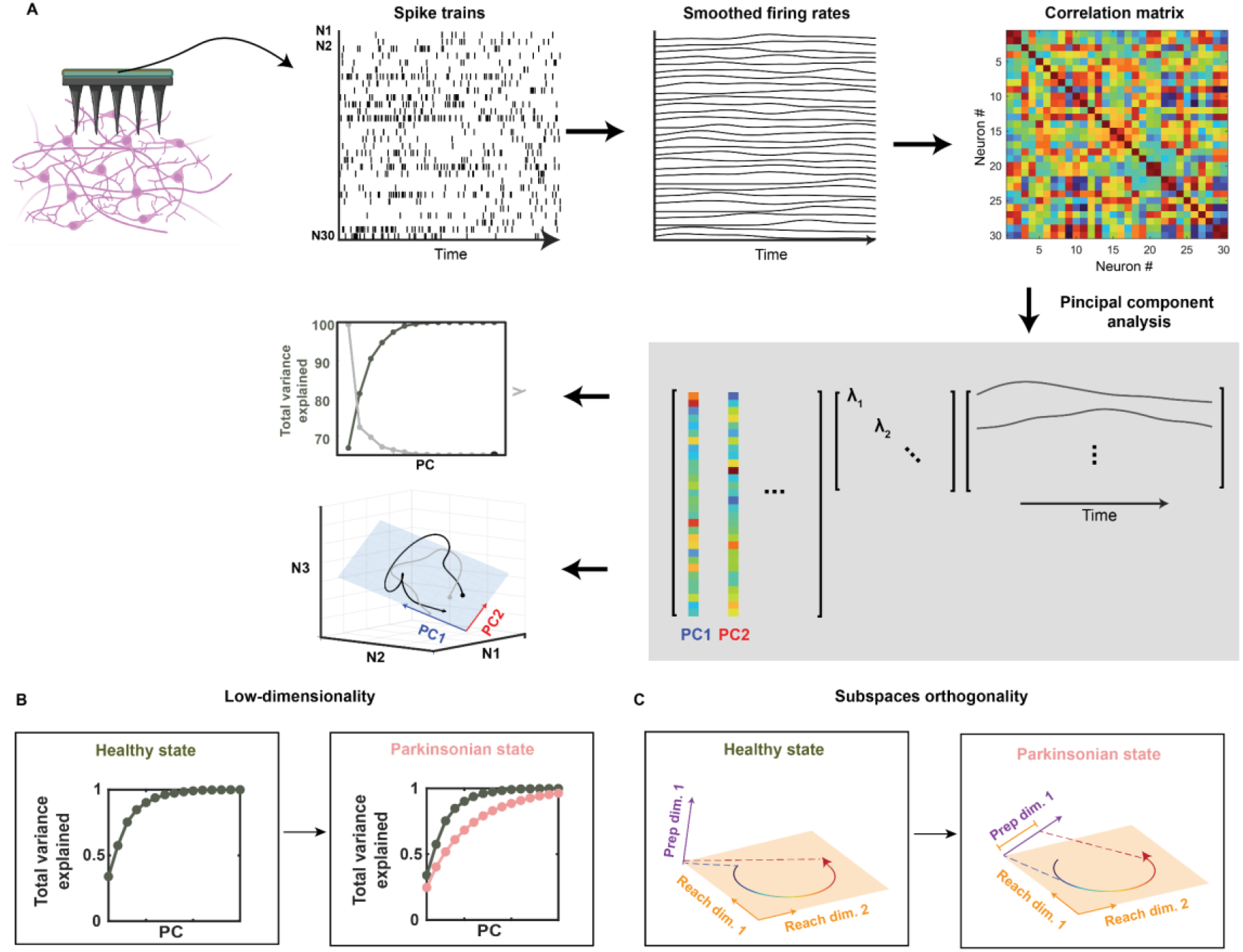
Putative population-level organizational changes. (A) Inferring population-level organization of cortical dynamics. Spike trains from simultaneously recorded neurons (n = 30, example subset shown) are smoothed to obtain continuous firing rates, from which pairwise correlations are computed. Principal component analysis (PCA) is then applied to identify dominant patterns of shared variability, yielding a set of population modes (principal components) characterized by neuron loadings and their corresponding temporal activity profiles. In this illustrative example, the first two principal components explain >80% of the total variance and define a two-dimensional subspace that captures the dominant structure of population activity. The activity of three example neurons (selected from the recorded population for visualization) is shown to illustrate how low-dimensional population dynamics capture single-neuron activity. (B) Cumulative variance explained by population activity as a function of the number of principal components (PCs) in the healthy (green) and parkinsonian (pink) states, illustrating differences in the effective dimensionality of neural activity during the task. (C) Schematic illustration of neural state trajectory during reach epoch. Axes represent the leading preparatory dimension (purple) and the first two reach-related dimensions (orange), identified by dimensionality reduction applied to population activity from N neurons. In the healthy state (left), the preparatory- and reach-related subspaces are approximately orthogonal, resulting in minimal projection of reach-epoch activity onto the preparatory-related subspace. In contrast, in the parkinsonian state (right), partial overlap between the two subspaces leads to increased projection of reach-epoch activity onto the preparatory-related subspace.

In this study, we simultaneously recorded populations of neurons in the primary motor cortex (M1) from two nonhuman primates performing reaching tasks before and after induction of parkinsonism with the neurotoxin 1-methyl-4-phenyl-1,2,3,6-tetrahydropyridine (MPTP). In the parkinsonian state, both movement preparation and reach durations were significantly prolonged. By applying dimensionality reduction, we found that parkinsonism altered the structure of M1 activity at the population level, manifesting as increased dimensionality during both the preparation and movement epochs and increased alignment between preparatory and reach-related subspaces. Furthermore, in the parkinsonian state, the trial-by-trial variability in reach duration was explained by the alignment between preparatory-related and reach-related subspaces, indicating the functional relevance of subspace orthogonality in PD. Together, these results motivate a population-level view of the motor dysfunction that occurs in PD where the flexibility in switching neural dynamics across computations is reduced in the parkinsonian condition, leading to degraded motor outcomes.

## Results

### Impaired movement preparation and execution in MPTP model of NHP during cued reaching task

Two non-human primates (NHPs) were trained on two reaching tasks in their naïve states: a reach-to-grasp task and a center-out touchscreen task (Figure 2A,B, see methods for detailed descriptions). We collected the kinematics data and neural signals from the 96-channels Utah arrays implanted in the primary motor cortex (M1) while the monkeys performed the tasks before and after the induction of parkinsonism with 1-methyl-4-phenyl-1,2,3,6-tetrahydropyridine (MPTP). After MPTP treatment, both monkeys exhibited moderate parkinsonism during the recording sessions as indicated by the mUPDRS scores (Table 1). During the cued reaching task, we found that both the movement preparation and execution were impaired in parkinsonian monkeys indicated by the prolonged preparation and reach durations (Figure 2C). In addition, preparation and reach durations showed greater variability in the parkinsonian state, consistent with clinical reports in PD patients^26–28^.

**Table 1.** Clinical ratings (mUPDRS) of parkinsonian motor sign.

| **Motor symptoms (maximum score)** | **Animal J** | **Animal K** |
| --- | --- | --- |
| **Rigidity (6):** | 1.6 ± 0.2 | 4.0 ± 0.2 |
| **Tremor (6):** | 0.0 ± 0.0 | 0.6 ± 0.0 |
| **Bradykinesia (6):** | 3.8 ± 0.2 | 6.0 ± 0.0 |
| **Akinesia (6):** | 3.8 ± 0.4 | 6.0 ± 0.0 |
| **Food Retrieval (3):** | 1.5 ± 0.0 | 2.5 ± 0.0 |
| **Axial (15)** | 8.2 ± 0.9 | 11.2 ± 0.6 |
| **Total\* (42):** | 18.9 | 30.3 |

Values are mean ± SD. Number of observations for both animals were 6, except for axial scores: 3 observations for animal J in parkinsonian state; 5 observations for animal K in parkinsonian state.

**Figure 2:**
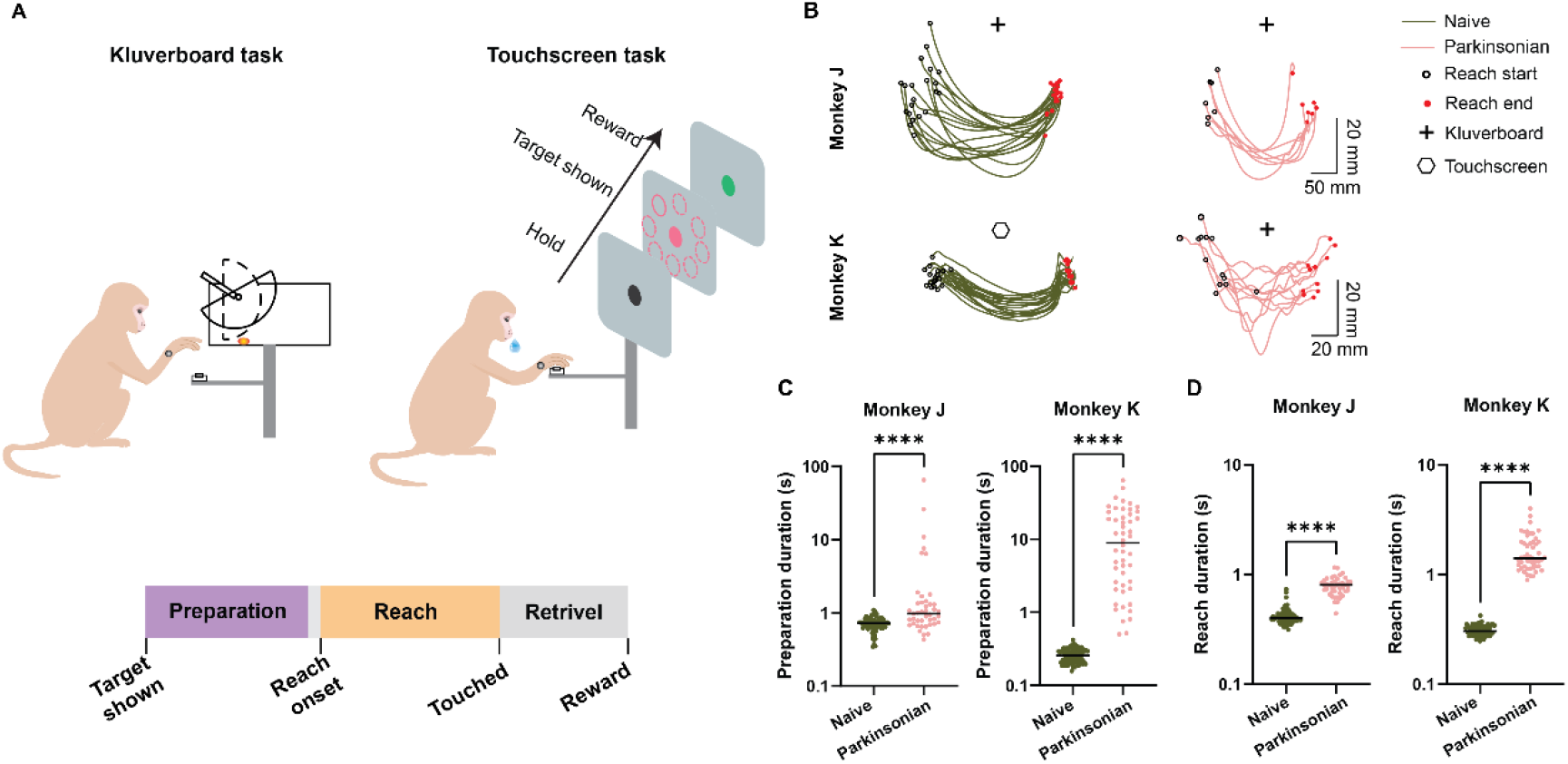
Behavioral tasks and performances before and after the induction of parkinsonism. (A) Illustration of the reaching tasks (top) and task timeline (bottom). Colored areas mark the preparatory (from target shown to 50 ms prior to the reach onset, defined by wrist speed; purple) and reach epoch (from reach onset to touched; orange) analyzed in this study. (B) Reaching trajectories projected onto the plane capturing the largest variability from example sessions, with line color coded for condition. (C) Preparation duration is increased in parkinsonian state compared to the naive state. (D) Reach duration is increased in parkinsonian state compared to the naive state. (C, D) Y-axis is plotted on a logarithmic scale. Wilcoxon rank sum test, p < 0.0001 for both Monkey J and Monkey K.

### Low-dimensional subspace is disrupted in parkinsonism

We recorded single-neuron responses from the primary motor cortex in the naive (240 neurons from monkey J across 3 sessions, 259 from monkey K across 3 sessions) and parkinsonian states (135 neurons from monkey J across 3 sessions, 278 from monkey K across 4 sessions). Consistent with the previous findings on naïve monkeys, single unit firing shows complex, heterogeneous responses during the reaching task^14,19^, with individual neurons preferentially responding to different task events during preparation or movement epochs (Supp Fig 1 upper). This heterogeneity of response was also observed in the parkinsonian states (Supp Fig 1 bottom). The parkinsonian condition was characterized by substantial trial-by-trial variability in preparatory and reach durations (Figure 2C, D). This variability resulted in pronounced temporal misalignment of neural activity relative to behavioral epochs across trials (Figure 3A), such that trial-averaging may obscure structured dynamics. Therefore, rather than imposing a categorical structure on heterogeneous and temporally variable single-neuron responses, we focused on population-level analyses that capture single-trial coordinated neural dynamics (see *Methods*).

**Figure 3:**
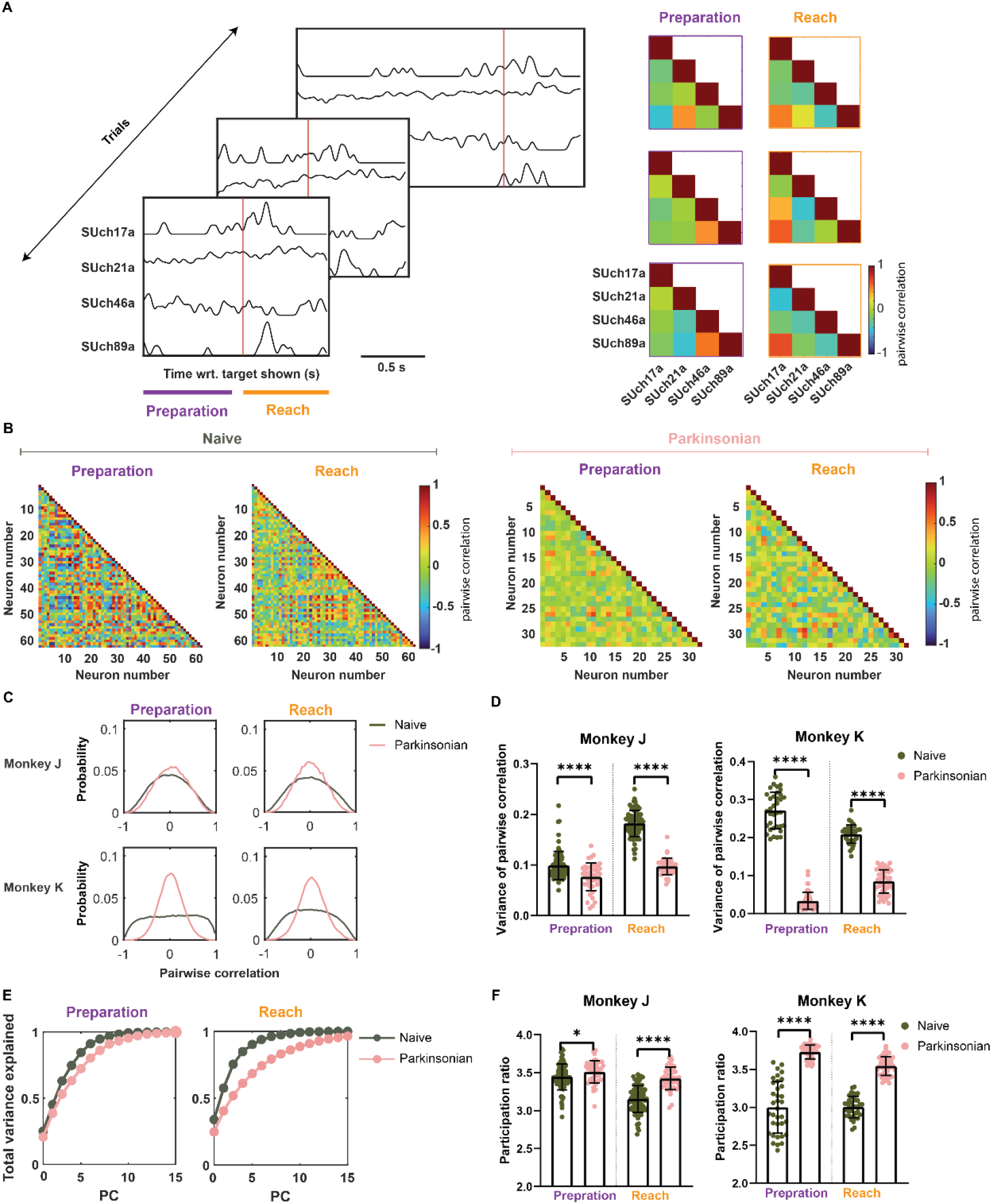
Neural coordination in naive and parkinsonian states. (A) Neuronal response from four example units aligned to the target shown across representative trials in parkinsonian state. Red vertical lines indicate reach onset. Preparatory (purple) and reach (orange) epochs are indicated by colored bars beneath the time axis. Trials are reordered by the combined duration of the preparatory and reach epochs. Individual neurons exhibit heterogeneous temporal response profiles and pronounced trial-by-trial variability in response timing. For each corresponding trial, pairwise correlation matrices (right) are computed from single-trial neuronal response separately within the preparatory and reach epochs, revealing epoch-dependent coordination across units. Each entry in the matrix reflects the similarity between the response patterns of the two neurons during that epoch. (B) Correlation matrices for all neurons during preparatory epoch and reach epoch of example trials from naive (left) and parkinsonian (right) states in Monkey J. (C) Distribution of pairwise correlations across all neuron pairs and trials, pooled across sessions within condition, during the preparatory epoch (left) and the reach epoch (right). (D) Trial-wise variance of the pairwise correlation distribution across neuron pairs during the preparatory and reach epochs for naive and parkinsonian states.

Previous studies have shown that neuronal activity at the population level is confined to low-dimensional subspaces spanned by a few independent patterns during the movement preparation and execution^17,19,22,25^. These low-dimensional subspaces are likely to emerge from the coordinated activity of interacting neurons. Here, an example pairwise correlation matrix from our dataset shows that, during both the preparatory and reach epochs, pairwise correlations in the naive state span a broader range, with more pronounced positive and negative correlations (Figure 3B, left) compared to parkinsonian condition (Figure 3B, right). When pairwise correlations were aggregated across all trials, the parkinsonian state exhibited a markedly narrower distribution than the naive state in both epochs, despite similar near-zero means (Figure 3C). This compression of correlation structure was further quantified as a significant reduction in the variance of pairwise correlations in the parkinsonian state during both the preparatory and reach epochs (Figure 3D), which suggests reduced coordination within the population in the parkinsonian state. This decorrelation would be expected to distribute variance across more dimensions. Indeed, in example trials, more principal components were required in the parkinsonian state to explain the same amount of variance as in the naive state (Figure 3E). To quantify this effect, we computed the participation ratio (PR) as a measure of effective dimensionality^29–31^. To control for biases arising from unequal number of recorded neurons and unequal epoch durations, we randomly subsampled neurons and data segments to match both neuron number and data length across the naive and parkinsonian states when computing the participation ratio. We found that the effective dimensionality in the parkinsonian state is higher than that in the naive state during both the preparatory and reach epochs (Figure 3F). To validate that our subsampling procedure did not bias the observed differences, we repeated the analysis shown in Figure 3F using all neurons recorded from Monkey K, which had a larger number of recorded neurons than Monkey J. The increase in dimensionality in parkinsonian state remained qualitatively unchanged (Suppl fig 2).

These variances were pooled across trials and sessions for visualization. Bars represent mean ± s.d. (E) Cumulative variance explained as a function of the number of principal components (PCs) during the preparatory (left) and reach (right) epochs from example trials of naïve and parkinsonian conditions in Monkey J. (F) Trial-wise participation ratio during the preparatory and reach epochs for naive and parkinsonian states. These participation ratios were pooled across trials and sessions for visualization. Bars represent mean ± s.d.

### Orthogonality between preparatory and reach subspaces is impaired in Parkinsonism

Having shown that parkinsonism alters the low-dimensional structure of population activity in M1 during movement preparation and execution, we next asked whether it also alters the relationship between the preparatory and reach subspaces. In healthy animals, these subspaces are nearly orthogonal and this orthogonality is suggested to provide a mechanism to segregate motor planning from execution, thereby being critical for motor control^19,20^. To quantify how strongly activity from each epoch occupied the two subspaces, we measured subspace alignment. We projected population activity from the preparatory and reach epochs onto the preparatory-related and reach-related subspaces identified from the corresponding epochs. For visualization, the trajectories corresponding to the first two dimensions of each subspace are shown. In the naïve state, preparatory activity evolves prominently within the preparatory-related subspace but exhibits minimal variance when projected onto the reach-related subspace. During the reach epoch, population activity expands extensively within the reach-related subspace while showing minimal structure in the preparatory-related subspace (Figure 4A). In the parkinsonian state, however, preparatory activity exhibits increased spread within the reach-related subspace, and reach-epoch activity shows substantial structure when projected onto the preparatory-related subspace, indicating greater overlap between preparatory and movement-related population dynamics. To quantify the alignment between the preparatory-related and reach-related subspaces, and to generalize the analysis to arbitrary dimensionality, we computed an alignment index^18,20^. This index is defined as the ratio between (i) the variance of neural activity during one epoch after projection onto the subspace defined by the other epoch and (ii) the variance of the same activity after projection onto its own subspace. When computing the alignment index, we defined the preparatory and reach subspaces using the first *n* and *m* principal components (PCs), respectively, where *n* and *m* correspond to the minimal effective dimensionalities observed during the preparatory and reach epochs across all trials in both naïve and parkinsonian states (n = 4, m = 4 for Monkey J; n = 3, m = 3 for Monkey K). This approach controls for biases arising from mismatched subspace dimensionalities while ensuring that each subspace captures the task-related covariance rather than noises. The alignment index of the parkinsonian state is significantly higher than that of the naive state (Figure 4B, two-sided nested t test, Monkey J: p = 0.0015; Monkey K: p = 0.0008), indicating that the preparatory subspace and the reach subspace are more aligned in the parkinsonian state. Furthermore, in the parkinsonian state, trial-by-trial reach duration increased with the alignment index, such that greater overlap between the preparatory and reach subspaces was associated with longer reach durations (Figure 4C). Together, these results demonstrate that parkinsonism increases the alignment between preparatory and reach subspaces, and that this altered population structure is functionally linked to impaired movement execution.

**Figure 4:**
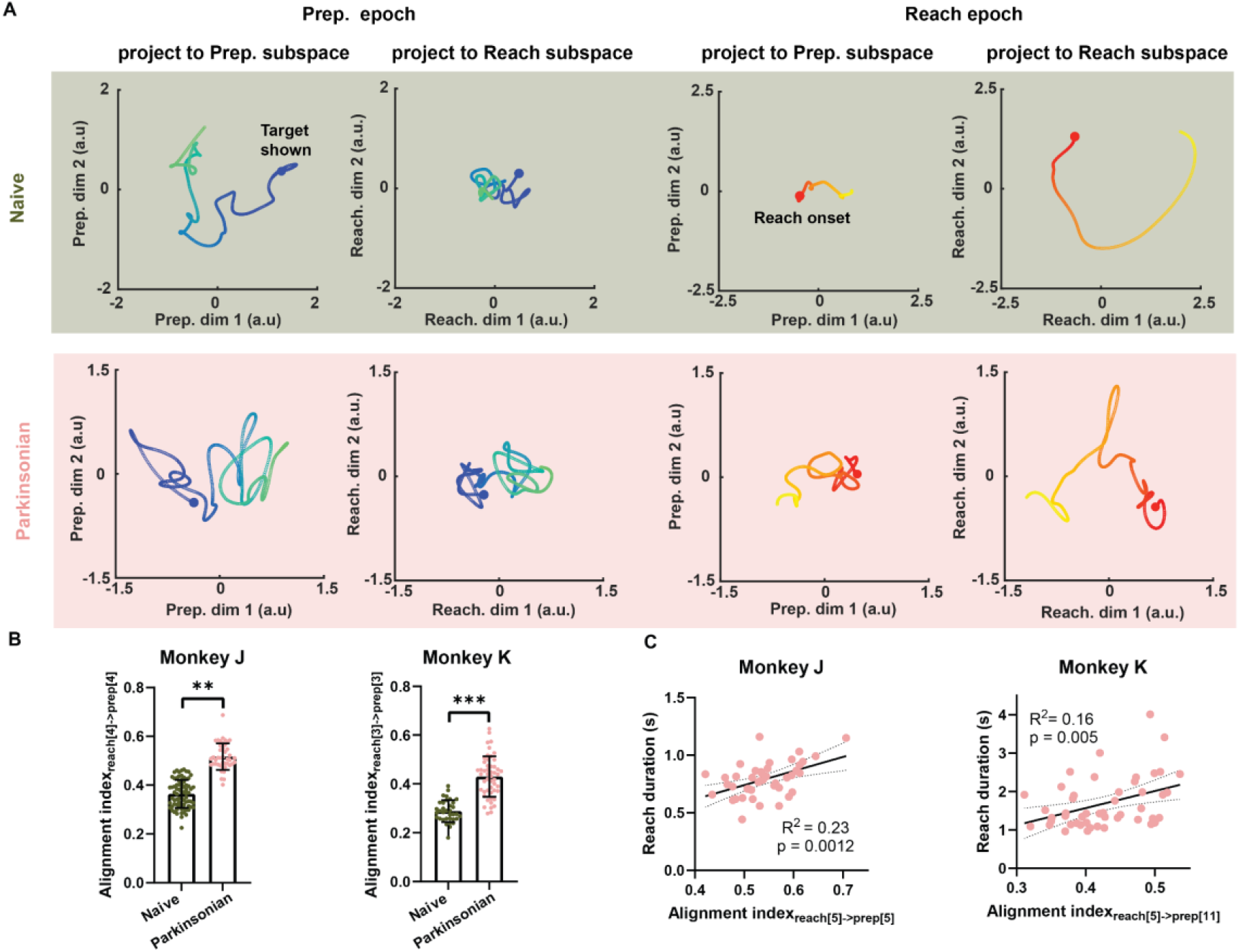
Reduced orthogonality between preparatory and reach subspaces. (A) Example of population responses during the preparatory (left two columns) and reach (right two columns) epochs in the naive (upper panel) and parkinsonian (bottom panel) states projected onto the first two dimensions of the preparatory-related subspace (column 1 and 3) and the reach-related subspace (column 2 and 4). Trajectories are color-coded by time, with task variables indicated (blue dot, target shown; red dot, reach onset). Data are from example trials in the naive and parkinsonian states in Monkey J. (B) Alignment index computed in the naive and the parkinsonian states. (C) Linear regression of reach duration in the parkinsonian state using alignment index. Axes are independently scaled for each subject for clarity. Solid line indicates the best linear fit; shaded area denotes the 95% confidence interval.

## Discussion

In this study, we show that parkinsonism disrupted the population-level organization of neural activity in the primary motor cortex during voluntary movement. By examining large-scale M1 recordings before and after MPTP-induced parkinsonism, we demonstrate that motor impairments are associated with increased neural dimensionality and reduced separation between preparatory and reach-related subspaces. By linking trial-by-trial variations in subspace orthogonality to prolonged reach durations, our findings suggest that reduced segregation between preparatory and movement-related dynamics contributes to impaired motor execution in parkinsonism. Together, these findings extend existing views of motor cortical dysfunction in Parkinson’s disease by framing parkinsonism as a disorganization of population-level neural dynamics.

Our findings highlight the importance of analyzing neural activity at the population level. Large-scale cortical recordings combined with dimensionality reduction approaches enable characterization of population structure that may constitute fundamental building blocks of neural computation but are not apparent at the level of individual neurons. Leveraging within-subject recordings in the naive and parkinsonian states, we further characterize how the structure is altered. Importantly, this experimental design and analytic approach allow us to uncover trial-by-trial relationships between neural subspaces and behavioral metrics. In much of the existing literature on Parkinson’s disease, neural metrics are reported to be altered or associated with parkinsonism; however, such representational changes alone do not establish functional relevance. By contrast, when variability in neural activity covaries with variability in behavior across trials, as quantified in our analyses, it supports the interpretation that these neural metrics are functionally relevant and more closely associated with the neural processes underlying action generation.

Low dimensionality is the empirical observation that, in most motor tasks, the number of neurons available to participate in a given cortical area far exceeds the number of independent variables required to specify the behavior. Such dimensionality arises from constraints on population activity and has been shown to scale with task complexity^23,32^. Neural activity therefore need not explore the full space of firing patterns, but instead varies along a limited set of directions that are relevant to task completion. Our findings suggest that parkinsonism may increase the effective computational demands of movement preparation and execution. Neural activity that is normally organized along a compact set of coordinated population modes during movement generation may become less structured in the parkinsonian state, potentially requiring additional corrective adjustments or engaging a broader set of neural degrees of freedom. This interpretation is consistent with the clinical reports of increased perceived difficulty during movement from PD patients^33^.

Beyond the increased dimensionality, we found that the parkinsonism disrupts the orthogonality between the population-level structure of the movement preparation and execution. Orthogonality is assumed to reflect a strategy of carrying out different computations (i.e., movement preparation and execution in our study) within the same population of neurons under active control. The association of reduced orthogonality between preparatory-related and reach-related subspaces and the impaired behavioral metrics found in our study supports the functional role of orthogonality. The question remains, however, concerning which neural circuit(s) gives rise to this orthogonality between preparatory and movement subspaces in the healthy state, how this orthogonality is established, and how parkinsonism disrupts this process. Kao et al. showed that an optimal feedback (rather than feedforward) preparatory control strategy can account for the observed orthogonality between preparatory and movement-related activity in monkey M1. This arises because feedback control selectively regulates only those state-space directions that have future motor consequences, while allowing other “motor-null” directions to vary without affecting the upcoming movement^34^. They further proposed that such preparatory feedback control could be implemented via a thalamo-cortical loop that is transiently engaged and gated by the basal ganglia. At the circuit-mechanistic level, Bachschmid-Romano et al. constructed a recurrent network model constrained by macaque M1 data and found that multiple combinations of recurrent connectivity and external inputs can reproduce the recorded dynamics^35^. Notably, among solutions, the minimum-input constraint assigns a distinct role to external inputs versus recurrent connectivity: during movement preparation, covariance structure is primarily shaped by direction-specific external inputs, whereas during movement execution it is dominated by strong direction-specific recurrent connectivity. This epoch-dependent shift in the balance between external input and recurrent dynamics lead to changes in single-neuron tuning across preparation and execution, which in turn can manifest as distinct population-level organization.

Parkinsonian pathophysiology encompasses multiple abnormalities that could perturb these mechanisms. Reduced thalamocortical synaptic efficacy and structural alterations at thalamocortical synapses have been reported in parkinsonian states^36,37^, and disrupted premotor–motor cortical connectivity has been observed, with abnormal PMd–M1 interactions partially normalized by dopaminergic therapy^38,39^. Such alterations in cortical inputs could compromise the efficacy of preparatory feedback control and reduce the reliability with which M1 is steered into movement-appropriate initial conditions. In addition, dopamine-dependent changes in cortical network organization, including reduced monoaminergic innervation ^40,41^ and elevated spine elimination and formation in the parkinsonian M1^42^, may further modify intrinsic connectivity and recurrent dynamics. Together, these abnormalities are consistent with a disruption of the mechanisms that normally maintain orthogonality between preparatory and movement-related activity, potentially contributing to the reduced subspace separation observed here.

An alternative, non–mutually exclusive explanation for the reduced subspace orthogonality in Parkinsonism involves the presence of excessive beta-band activity, a prominent feature of the parkinsonian motor system^43,44^. If pathological beta activity persists across both preparatory and execution epochs, it could impose a shared temporal modulation on neuronal firing that constrains the reconfiguration of population activity across task phases. Under such conditions, preparatory and execution activity may be biased toward overlapping dimensions, thereby reducing the apparent orthogonality between the corresponding subspaces. Consistent with this possibility, recent work demonstrated that impaired motor performance is associated with a loss of temporal separability between LFP beta bursts and movement-related ensemble co-firing quantified using dimensionality reduction^45^, whereas behavioral recovery coincided with a restoration of this separability. These findings suggest that pathological beta activity may intrude into neural states normally associated with movement execution, limiting the dynamic separation between task epochs. Similar considerations may also apply to other pathological activity patterns observed in PD, such as exaggerated phase–amplitude coupling (PAC)^46,47^, high-frequency oscillations (HFO)^48^ and aberrant bursting^49^, which could likewise impose constraints on population dynamics. Notably, excessive beta activity in Parkinson’s disease is most robustly observed in subcortical structures of the basal ganglia–thalamocortical network^43,44^, whereas cortical beta abnormalities, particularly in primary motor cortex, are less consistent; indeed, in our prior work, we did not observe a consistent elevation of beta power in M1 at rest^50^. Together with the fact that beta oscillations are typically characterized at the level of local field potentials, these observations indicate that any influence of pathological beta on cortical population structure is likely to be indirect or task-dependent. Thus, while excessive beta activity provides a plausible population-level constraint that could contribute to reduced preparatory–execution orthogonality, establishing a direct relationship between beta dynamics and subspace geometry will require explicit joint analyses of spiking activity and field potentials during active movement tasks in the future.

The movement-related population-level changes at the cortical level reported here were consistent across animals. This contrasts with our prior findings from resting-state cortical recordings, which revealed heterogeneous changes in beta oscillatory power across animals^50,51^. Similarly, previous studies have reported inconsistent effects of therapeutic interventions such as deep brain stimulation and levodopa on cortical beta^52,53^. Together, these observations indicate the importance and additional value of investigating movement-related neural activity in order to fully understand the pathophysiology related to the functional changes in PD.

Our findings open several additional avenues for future investigation. In particular, it will be important to determine how the basal ganglia–thalamo–cortical network contributes to preparatory–movement subspace orthogonality in the healthy state and how these circuit mechanisms are altered in parkinsonism. An additional key question is whether clinically effective therapies, such as levodopa and deep brain stimulation, restore the cortical population-level organization disrupted in PD. Together, these findings highlight population-level organization as a framework for understanding motor dysfunction in Parkinson’s disease and may help guide the development of novel biomarkers and therapeutic strategies for its treatment.

## Methods

### Experimental subjects and procedures

Two adult female rhesus macaques (Macaca mulatta), aged 17 (Monkey J) and 13 (Monkey K), were included in this study. After the initial training on reaching tasks (described below), each monkey was implanted with a 96-channel Utah microelectrode array (Pt-Ir, 1.5 mm depth, 400 um interelectrode spacing, Blackrock Microsystems) in the arm area of the M1. The instrumentation and surgery details have been described previously^50^. All procedures were approved by the University of Minnesota Institutional Animal Care and Use Committee and complied with US Public Health Service policy on the humane care and use of laboratory animals.

### Induction of parkinsonism

Following normal state recordings, animals were rendered parkinsonian through intra-carotid and systemic intramuscular injections of 1-methyl-4-phenyl-1,2,3,6-tetrahydropyridine (MPTP). Monkey J received one intra-carotid injection and three intramuscular injections (0.3-0.4 mg/kg each, totaling 1.4 mg/kg), while Monkey K received six intramuscular injections (0.3-0.4 mg/kg each, totaling 1.8 mg/kg). Parkinsonian severity was evaluated using a modified Unified Parkinson’s Disease Rating Scale (mUPDRS), assessing axial motor symptoms (gait, posture, balance, turning, defense reactions), food retrieval, and appendicular motor symptoms (rigidity, bradykinesia, akinesia, and tremor) on the contralateral hemi-body, using a 0-3 scale (0 = normal, 3 = severe; max score = 42). Data collection occurred after both animals reached a moderately parkinsonian state, at least two months post-injection when the mUPDRS score was stable between 18 and 31^50^.

### Behavioral task and analysis

Monkey J was trained to perform a reach-to-grasp task, beginning with the hand on a start pad and a food reward presented in front of the monkey as a “go” cue. Monkey K performed the same reach-to-grasp task in the parkinsonian state as Monkey J, but a center-out touchscreen task in the normal state. In the center-out task, the monkey reached for different targets on the screen from the start pad after the target was shown. We only include centered target, successful trials in subsequent analyses to maintain consistency with the direction of food reward in the reach-to-grasp task. Overall, the reach-to-grasp task and the center-out task are similar in the reaching epoch and commonly used to study the neural mechanism of reaching^17,18^.

The preparatory epoch is defined as the period from target shown to 50 ms before the movement onset. Reach epoch is defined as the period from movement onset to the target touched. We defined reaction time and reach duration, the primary behavioral metrics, as the time spanning preparatory and reach epoch. Kinematic data were recorded with a passive marker motion capture system (Motion Analysis) at 100 Hz, tracking 3D wrist, elbow and shoulder positions. Wrist speed was calculated by differentiating the smoothed (cubic spline) wrist position data. Reach onset and end were determined when the wrist marker’s speed exceeded 10 mm/s.

### Neuronal data recording and analysis

Neurophysiological data were recorded using a TDT workstation (Tucker Davis Technologies). Signals were band-pass filtered (300 Hz–5 kHz) and single units were isolated and sorted using principal component and template-based methods in Offline Sorter (Plexon, USA). To obtain the continuous firing profiles of recorded units, we binned the spikes with 1 ms window and smoothed with a Gaussian kernel (*σ* = 20 ms). To ensure reliable estimation of population activity structure, we excluded units with mean firing rates below 5 Hz during the analytical window.

### Population analysis via PCA

For analysis of dimensionality and subspace identification, we applied PCA to data matrix *X* ∈ ℝ^*T×N*^ of continuous firing rates of *N* units in a *T* time window after soft normalization and mean-centering described in Churchland et al.^14^. PCA will decompose the covariance matrix of *X* as *X*^*T*^*X=VΣ*^2^*V*^*T*^, where *Σ*^2^ is a diagonal matrix whose diagonal entries are the eigenvalues *λ*_1_, *λ*_2_, . . ., *λ*_*N*_ of the covariance matrix, and *V* is the eigenvector matrix which yielded *N* PCs.

### Estimation of neural dimensionality

We used participation ratio (PR) to measure the neural dimensionality^30^. PR counts the effective dimensions along which data are spread as a ratio of the square of the first moment and the second moment of the eigenvalue probability density function. Specifically,

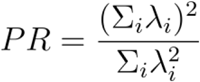

PR is thus in a range of 1 to N, with N being the total number of units. To ensure that PR was comparable across recording sessions with different numbers of neurons and varying epoch durations, we implemented a random downsampling procedure. For each trial and for each of 1,000 repetitions, neurons were randomly sampled to match the minimum number of recorded units (N = 25), while a consecutive 200 ms time window was randomly selected within the epoch of interest. PR was computed for each subsample. The mean PR across iterations was used for subsequent comparisons and statistical analyses.

### Subspace identification and alignment index

From the *N* PCs obtained by performing PCA on the data from the preparatory epoch, we defined *m*-dimensional, trial-specific preparatory-related by keeping only the leading *m* PCs. Similarly, we can obtain *k*-dimensional reach-related subspace. *m* and *k* are determined as the effective dimensionality of corresponding epoch. For analyses involving comparisons across conditions, *m* and *k* were matched across conditions by selecting the minimum effective dimensionality observed across trials, sessions, and conditions. For analysis restricted to the parkinsonian condition, *m* and *k* were determined using the minimum effective dimensionality across trials and sessions within parkinsonian condition only.

To quantify the amount of variance shared between the preparatory and movement population responses, preparatory-epoch activity was projected onto the reach-related subspace (or vice versa), and the total variance captured by this cross-projection was computed (“across epoch variance explained”). For normalization, activity from each epoch was also projected onto its own subspace, and the total variance captured was computed (“within-epoch variance explained”). The alignment index was defined as the ratio of across-epoch variance explained to within-epoch variance explained. Though the alignment index generally reflects the similarity of two subspaces, this measure is directional, which can be formally written as:

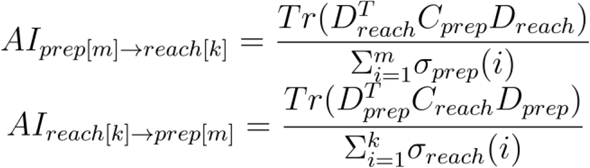

Where *C* is the covariance matrix of the data matrix *X*. *D*_*prep*_ is the matrix defined by the top m prep-PCs while *D*_*reach*_ is the matrix defined by the top k reach-PCs. *σ*(*i*) is the th singular value of *C*.

### Statistics

Behavior metrics (preparation and reach durations) between the naive and parkinsonian states were assessed using Wilcoxon rank sum test after pooling trials from multiple sessions within the condition.

Variance of pairwise correlation and participation ratio were computed on a trial-by-trial basis. Condition (Naive vs Parkinsonian) and epoch (preparatory vs reach) were treated as categorical fixed effects. To account for repeated measurements across sessions, we fit linear mixed-effects models separately for each monkey, with session included as a random effect. Planned contrasts were used to compare naive and parkinsonian states within each epoch based on the fitted model.

The alignment index between the naive and parkinsonian states were assessed using the nested t tests.

## Acknowledgements

We would like to thank our colleagues in the Neuromodulation Research Center for their helpful feedback on this study and especially thank our animal care team of Claudia Hendrix, Hannah Baker, and Elizabeth McDuell. Funding: This work was supported by the NIH, National Institute of Neurological Disorders and Stroke (R01-NS110613, R01-NS117822, R01-NS058945, R01-NS037019, R37-NS077657, P50-NS123109), American Parkinson Disease Association (Post-Doctoral Research Fellowships), Minnesota’s Discovery Research and Innovation Economy Brain Conditions Program, and the Engdahl Family Foundation.

## Author contributions

L.A.J., J.W., and J.L.V. designed the experiment; L.A.J. collected the data; B.M. curated the dataset; Y.N. designed and performed the analysis; Y.N. wrote the paper; Y.N., L.A.J., J.W., H.S. and J.L.V. edited the paper.

## Supplementary figures

**Figure S1:**
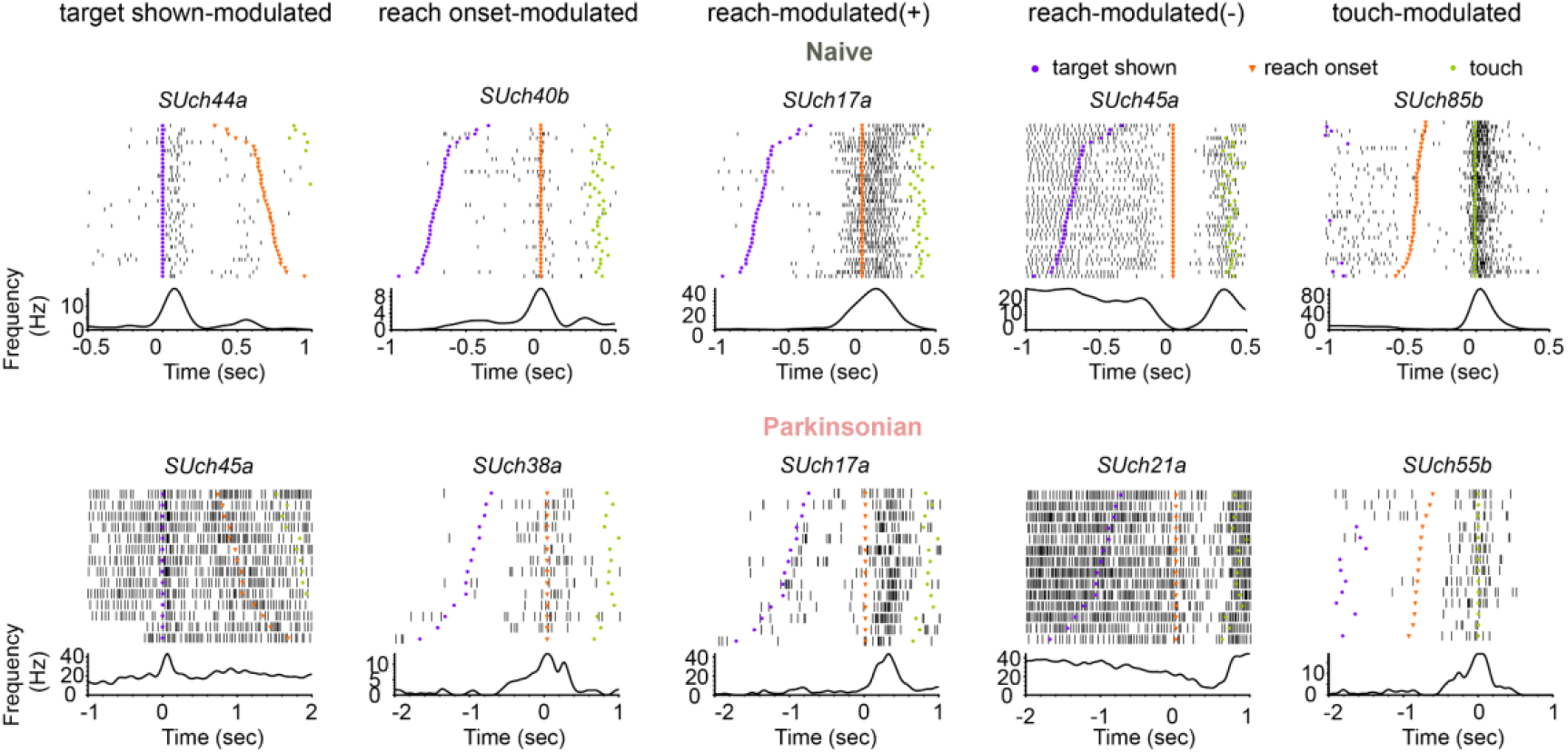
Raster plots and peri-event time histograms (PETHs) of example single-units of five categories in naive (upper) and parkinsonian (bottom) states. The preparatory epoch was defined as the interval from target shown (purple) to 50 ms prior to reach onset (orange). The reach epoch was defined from reach onset to touch. Target-shown–modulated neurons were modulated following target presentation (purple). Reach-onset–modulated neurons exhibited transient responses aligned to the reach onset (orange). Reach-modulated (+) and (−) neurons showed sustained increases or decreases in activity during the reach epoch, respectively. Touch-modulated neurons showed activity changes time-locked to target being touched (green).

**Figure S2.**
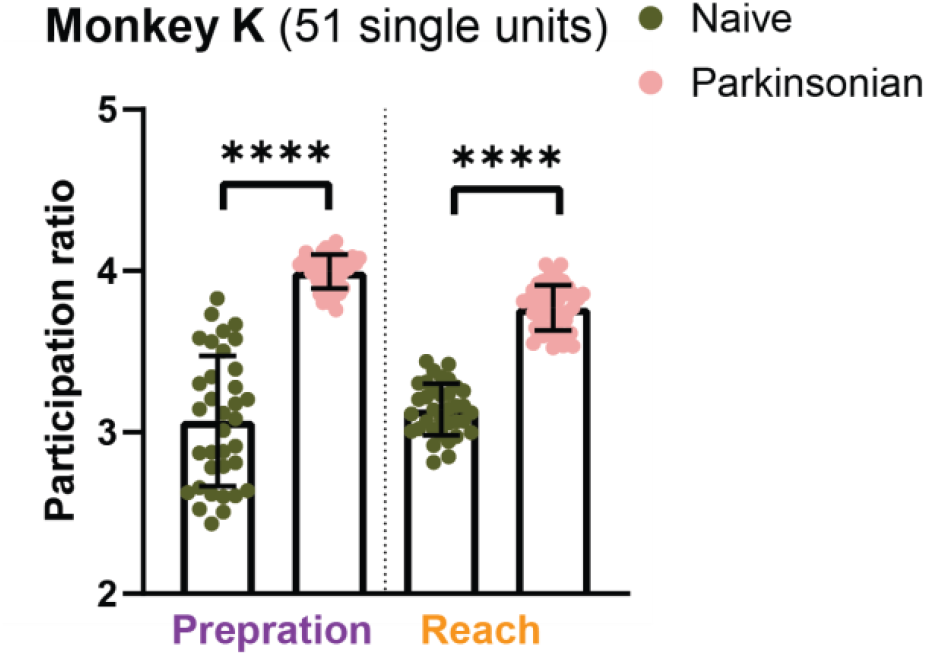
Participation ratio from Monkey K with larger number of single-units used, related to Figure 4. Trial-wise participation ratio during the preparatory and reach epochs for naive and parkinsonian states. These participation ratios were pooled across trials and sessions for visualization. Bars represent mean ± s.d.

